# Senescence-associated metabolomic phenotype in primary and iPSC-derived mesenchymal stromal cells

**DOI:** 10.1101/542357

**Authors:** Eduardo Fernandez-Rebollo, Julia Franzen, Jonathan Hollmann, Alina Ostrowska, Matteo Oliverio, Torsten Sieben, Björn Rath, Jan-Wilhelm Kornfeld, Wolfgang Wagner

## Abstract

Long-term culture of primary cells is reflected by functional and secretory changes, which ultimately result in replicative senescence. In contrast, induced pluripotent stem cells (iPSCs) do not reveal any signs of cellular aging while in the pluripotency state, whereas little is known how they senesce upon differentiation. Furthermore, it is largely unclear how the metabolome of cells changes during replicative senescence and if such changes are consistent across different cell types. In this study, we have directly compared culture expansion of primary mesenchymal stromal cells (MSCs) and iPSC-derived MSCs (iMSCs) until they reached growth arrest after a mean of 21 and 17 cumulative population doublings, respectively. Both cell types acquired similar changes in morphology, *in vitro* differentiation potential, up-regulation of senescence-associated beta-galactosidase, and senescence-associated DNA methylation changes. Furthermore, MSCs and iMSCs revealed overlapping gene expression changes during culture expansion, particularly in functional categories related to metabolic processes. We subsequently compared the metabolome of MSCs and iMSCs at early and senescent passages and observed various significant and overlapping senescence-associated changes in both cell types, including down-regulation of nicotinamide ribonucleotide and up-regulation of orotic acid. Replicative senescence of both cell types was consistently reflected by the metabolic switch from oxidative to glycolytic pathways. Taken together, long-term culture of iPSC-derived MSCs evokes very similar molecular and functional changes as observed in primary MSCs. Replicative senescence is associated with a highly reproducible senescence-associated metabolomics phenotype, which may be used to monitor the state of cellular aging.

## Introduction

*In vitro* culture of primary cells is associated with continuous changes that ultimately result in replicative senescence: The proliferation rate declines, cells enlarge, and they lose differentiation potential (Campisi & d’Adda di Fagagna, 2007). These profound changes in the course of culture expansion hamper reproducibility of experiments, which is of particular relevance in regenerative medicine (Wagner & Ho, 2007). For example, mesenchymal stromal cells (MSCs) acquire continuous changes in gene expression and DNA methylation over subsequent passages, which can be used to track the state of cellular aging (Koch et al., 2012; Li et al., 2017; Schellenberg et al., 2014; Wagner et al., 2008). In contrast to quiescent or apoptotic cells, senescent cells are highly metabolically active (Wiley & Campisi, 2016). Senescent cells secrete a characteristic cocktail of interleukins, chemokines, growth and inflammatory factors, which compose the senescence-associated secretory phenotype (SASP) (Coppé et al., 2008). The SASP was meanwhile observed across multiple cell types and there is evidence that this secretory function of senescent cells impacts on wound healing, embryonic development, and tumorigenesis (Watanabe, Kawamoto, Ohtani, & Hara, 2017). However, little is known about the metabolomic changes that may be associated or even contribute to the process of cellular aging.

In contrast to primary cells, such as mesenchymal stromal cells, induced pluripotent stem cells (iPSCs) do not reveal any signs of replicative senescence (Koch et al., 2013; Lapasset et al., 2011). They can be culture expanded for many passages without reduced proliferation, loss of differentiation potential, or telomere attrition. In fact, upon reprogramming, iPSCs even seem to further converge towards a primitive ground state with serial passage (Hackett et al., 2013). It is unclear how iPSCs escape from replicative senescence, while there is evidence that their progeny is bound to this destiny again upon exit from pluripotent state (Frobel et al., 2014). Long-term growth curves and the molecular sequel of replicative senescence have so far hardly been systematically addressed in iPSC-derived cells. A better understanding of cellular aging upon differentiation of iPSCs is therefore urgently needed with regard to the enormous hopes for regenerative medicine.

Various protocols have been described to generate iPSC-derived MSC-like cells (iMSCs) (Diederichs & Tuan, 2014; Frobel et al., 2014; Kang et al., 2015). These cells closely resemble their primary counterpart in morphology, immunophenotype and three-lineage differentiation potential toward osteocytes, chondrocytes and adipocytes (Sabapathy & Kumar, 2016). It has been suggested that iMSCs might be more homogeneous than primary MSCs, which are well known to comprise multiple subpopulations (Ho, Wagner, & Franke, 2008). However, on the epigenetic level MSCs and iMSCs still remained distinct and DNA methylation patterns that reflect the tissue of origin or donor age were not recapitulated in iMSCs (Frobel et al., 2014). So far, it was unclear whether iMSCs undergo the same molecular changes as primary MSCs during culture expansion. In this study, we have therefore compared functional, transcriptomic and metabolomic changes during long-term growth of MSCs and iMSCs.

## Results

### iPSC-derived mesenchymal stromal cells are bound to replicative senescence

Mesenchymal stromal cells at first passage (n = 5) were reprogrammed into iPSCs and then re-differentiated toward MSCs (iMSCs). Syngenic MSCs and iMSCs were subsequently expanded until the cells entered proliferation arrest. MSCs proliferation rates decreased after about 20 to 40 days, whereas iMSCs proliferated slower with a later decline in proliferation rate. Within three months all cell populations reached the senescent state with a mean number of population doublings of 21.3 ± 1.4 and 17.1 ± 3.8 for MSCs and iMSCs respectively (Fig. 1A). The changes in cellular morphology were very comparable between MSCs and iMSCs: At early passages they displayed spindle shaped fibroblast like morphology, whereas cells at later passage were enlarged with flattened “fried egg” morphology (Fig. 1B).

**Figure 1:**
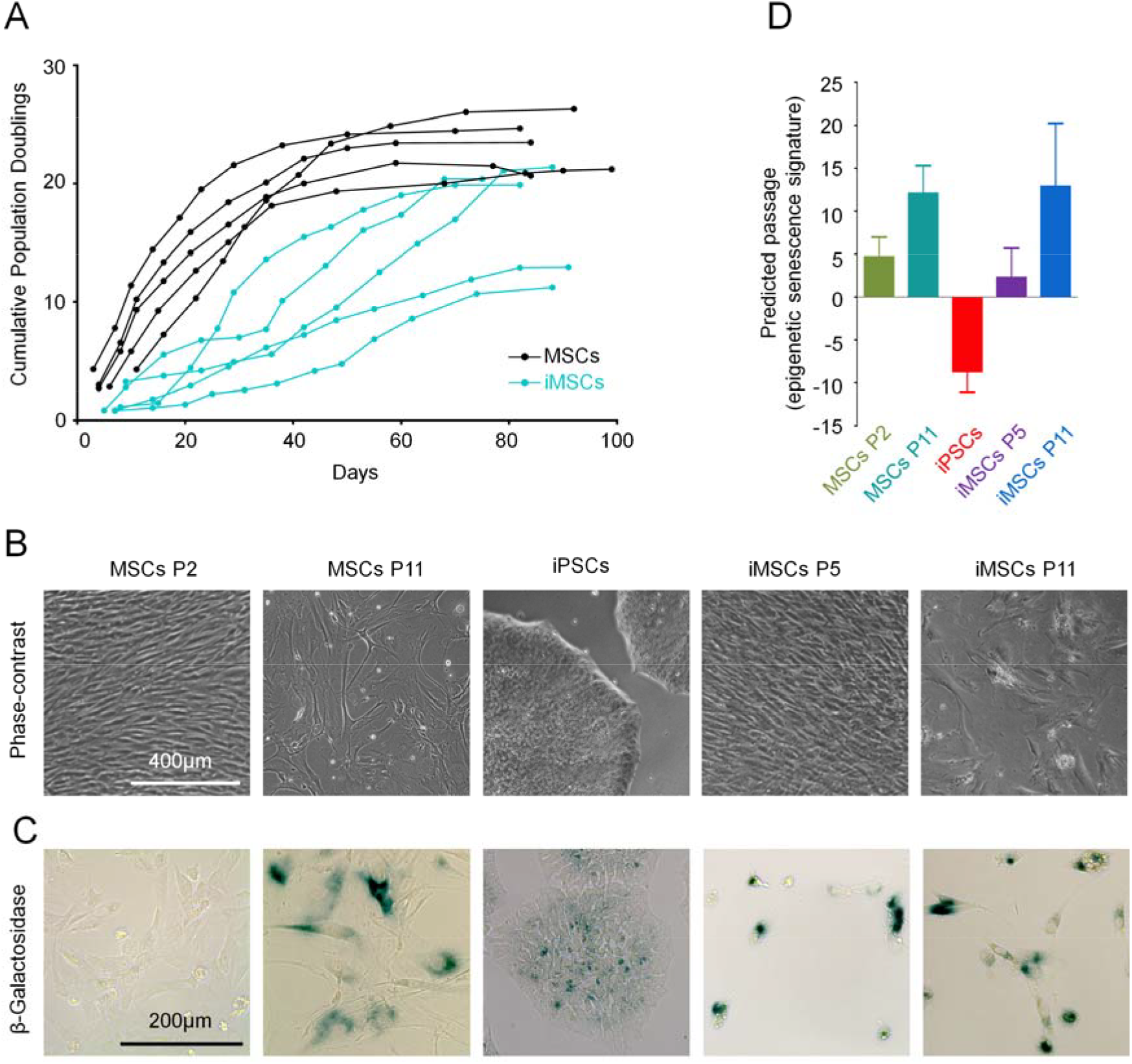
Replicative senescence of iPSC-derived MSCs. (A) Long-term growth curves of syngenic MSCs and iMSCs demonstrate that both cell types enter proliferative arrest within three months. (B) Phase contrast images of MSCs (passage 2 and 11) and iMSCs (passage 5 and 11). (C) Staining for senescence-associated beta-galatosidase (SA-β-gal) was more pronounced at later passages, albeit several iPSCs and iMSCs at early passage were also detected positive. (D) The number of passages was estimated based on an epigenetic senescence signature that utilizes DNA methylation levels at six CG dinucleotides (Koch et al., 2012).

The immunophenotype of MSCs and iMSCs was similar (CD29+, CD73+, CD90+ CD105+, CD14-, CD31-, CD34- and CD45-) and remained stable throughout culturing (Supplemental Fig. S1A). MSCs and iMSCs of early passages could be differentiated *in vitro* toward osteogenic, adipogenic and chondrogenic lineage (Frobel et al., 2014). The differentiation potential of MSCs decayed at later passages, as described before (Wagner et al., 2008), and this was also observed for iMSCs (Supplemental Fig. S1B).

Both cell types revealed increased staining for senescence-associated beta-galactosidase (SA-β-gal) at later passages, albeit several iPSCs and MSCs at early passage also stained positive for SA-β-gal (Fig. 1C). Subsequently, we estimated the state of cellular aging based on epigenetic modifications. We previously demonstrated that the number of passages is reflected by concurring DNA methylation changes at six senescence-associated CG dinucleotides (Franzen et al., 2017; Koch et al., 2012). MSCs and iMSCs revealed increasing epigenetic senescence predictions at later passages, whereas iPSCs were even predicted to be of negative passages (Fig. 1D). Taken together, iMSCs fulfilled the minimal criteria for definition of MSCs (Dominici et al., 2006) and both cell types apparently reveal very similar modifications during long-term culture that ultimately result in replicative senescence.

### Senescence-associated gene expression is closely related in MSCs and iMSCs

It has been demonstrated that long-term culture of MSCs is reflected by unique changes in their gene expression profiles (Wagner et al., 2008). To address the question if iMSCs reflect similar modifications we compared the transcriptome of MSCs and iMSCs at early and late passage by deep sequencing (n = 5). Overall the gene expression profiles of MSCs and iMSCs were closely related, albeit they were clearly separated by hierarchical clustering regarding to cell types and passages (Fig. 2A).

**Figure 2:**
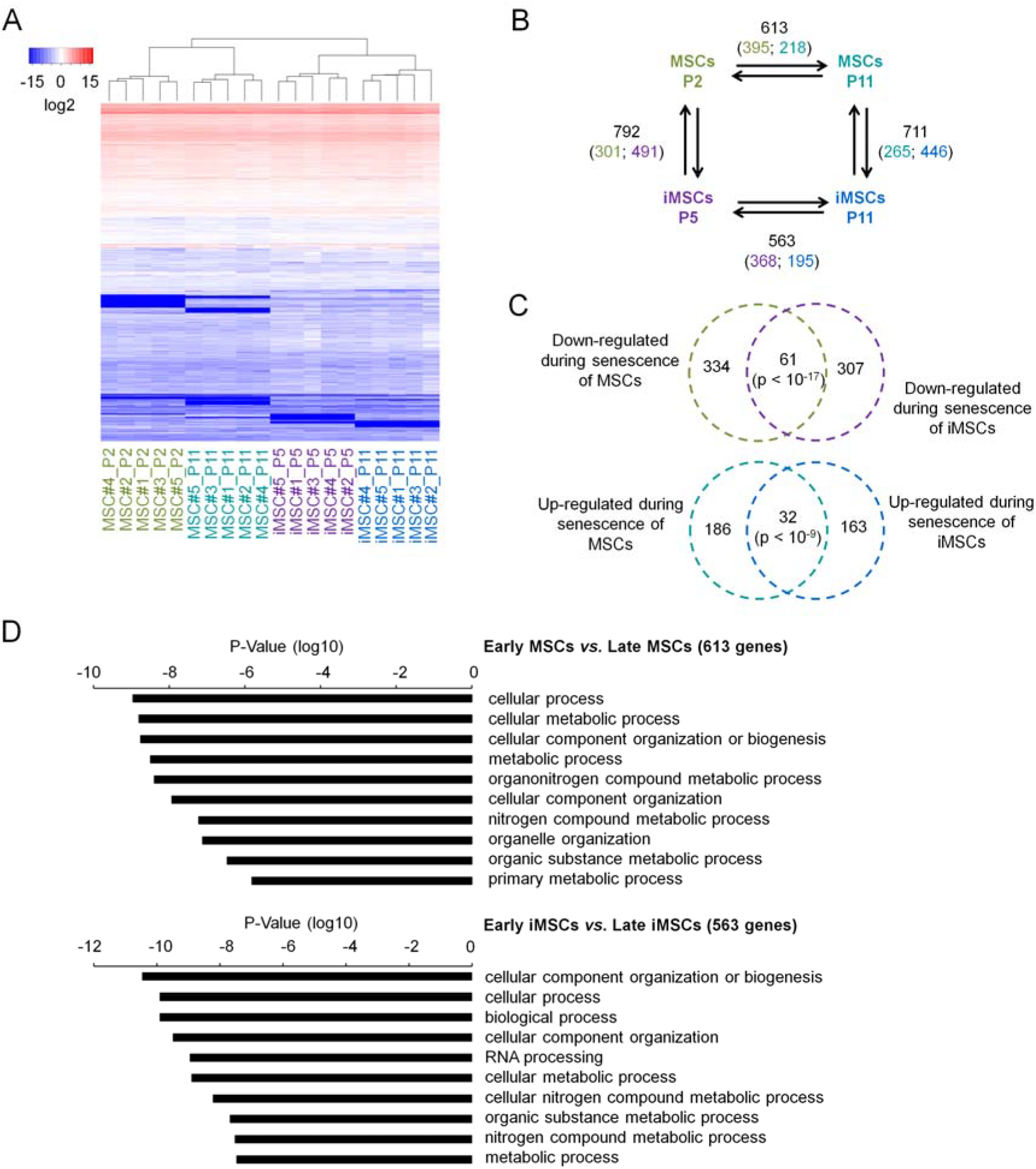
Transcriptome analysis of replicative senescence. (A) Heatmap of the entire transcriptome of MSCs and corresponding iMSCs (n = 5, donor numbers are indicated). Hierarchical clustering clearly separated early and late passages for both cell types. (B) The numbers of transcripts with significant gene expression changes in pairwise comparisons are depicted (2-fold differential expression and limma adjusted p value <0.05; the number of up-regulated genes in the corresponding cell type is indicated by color code). (C) The Venn diagrams depict significant overlaps of down- and up-regulated transcripts during senescence of MSCs and iMSCs (significance was estimated by hypergeometric distribution). (D) The senescence-associated gene expression changes in MSCs and iMSCs were classified by Gene Ontology analysis. Functional categories involved in metabolic processes were amongst the most significantly overrepresented (Fischer T-Test).

Pairwise comparison of MSCs and iMSCs revealed significant differences between early and late passages (2-fold differential expression and adjusted p value <0.05): 613 and 563 transcripts were differentially expressed upon senescence of MSCs and iMSCs, respectively (Fig. 2B). Notably, senescence-associated changes in MSCs and iMSCs share a relatively high number of down-regulated transcripts (61 transcripts; hypergeometric distribution: p < 10^−17^) and up-regulated transcripts (32 transcripts; p < 10^−9^; Fig. 2C). Senescence-associated gene expression changes were particularly enriched in gene ontology (GO) categories related to metabolism – such as cellular metabolic process, nitrogen compound metabolic process or organic substance metabolic process among others (Fig. 2D). This indicates that metabolic changes might be of particular relevance for the process of cellular aging in MSCs and iMSCs.

### Senescence is reflected by consistent changes in the metabolome

To gain insight into how the metabolism varies during long-term culture, we analyzed the metabolome of MSCs at passage 2 (n = 5) and passage 11 (n = 5) and of iMSCs at passage 5 (n = 5) and passage 11 (n = 5). For comparison we have also analyzed the five corresponding iPSC preparations. Using chromatographic approaches (UHPLC/MS/MS and GC/MS), 612 different metabolites were identified and measured. Hierarchical clustering of metabolomic profiles clearly separated cell types as well as early and late passages. In fact, MSCs and iMSCs at passage 11 comprised similar metabolites (Figure 3A).

**Figure 3:**
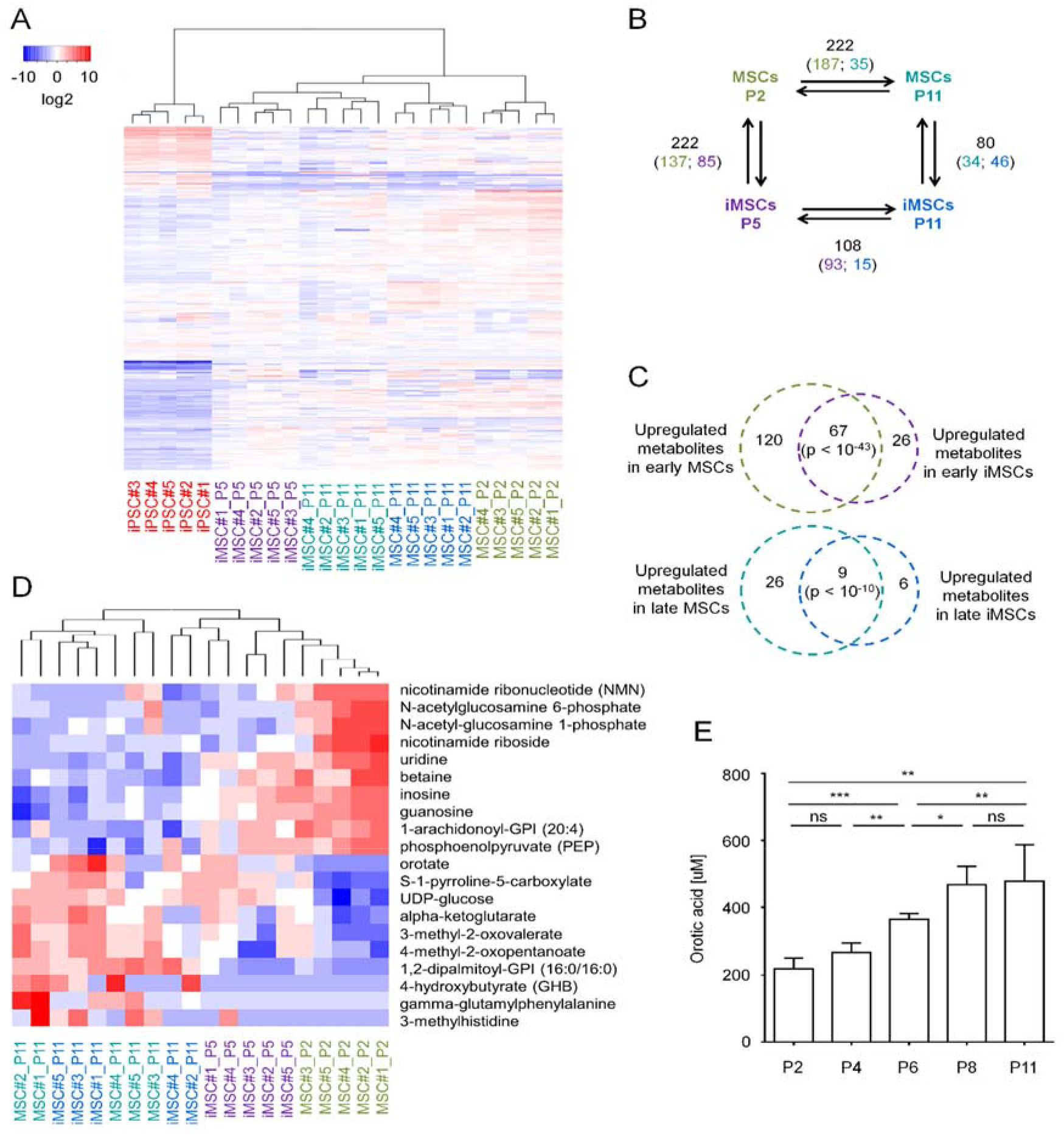
Metabolomic characterization of replicative senescence. (A) Heatmap presentation of metabolomes (612 different metabolites). Hierarchical clustering clearly separated iPSCs, MSCs, and iMSCs, as well as cells at early and late passage (n = 5, donor numbers are indicated). (B) Schematic presentation of the numbers of metabolites with significant differences in pairwise comparisons (1.5-fold difference and adjusted p value <0.05; higher abundance is indicated by the color code of corresponding cell type). (C) The Venn diagrams depict a high overlap of senescence-associated changes in MSCs and iMSCs for up-regulated and down-regulated metabolites (significance was estimated by hypergeometric distribution). (D) Heatmap of the 10 most significant up- and down-regulated metabolites during replicative senescence (MSCs + iMSCs). (E) Orotic acid was exemplarily measured by a fluorogenic assay in MSCs (n = 4) and increased over subsequent passages, which is in line with our metabolomics results.

We subsequently analyzed if there are significant differences in the metabolomic profiles of cells at early and late passage (>1.5-fold difference and adjusted p value <0.05): 222 and 108 metabolites were significantly changed in MSCs and iMSCs, respectively (Figure 3B). Notably, there was a highly significant overlap of senescence-associated metabolic changes in MSCs and iMSCs: 67 metabolites were consistently down-regulated (hypergeometric distribution: p < 10^−43^) and 9 metabolites were up-regulated at later passage in both cell types (p < 10^−10^; Figure 3C). In contrast, only sphingomyelin was significantly up-regulated during senescence of MSCs and down-regulated in iMSCs while inosine and uridine metabolites were up-regulated during senescence of iMSCs and down-regulated in MSCs. These results demonstrate that long-term culture of MSCs and iMSCs is overall associated with similar metabolomic changes. Amongst the metabolites with the most drastic down-regulation in senescence were N-acetylglucosamine 6-phosphate, N-acetylglucosamine 1-phosphate, nicotinamide riboside and nicotinamide ribonucleotide, whereas up-regulated metabolites included orotate, gamma-glutamylphenylalanine, 4-hydroxybutyrate and 3 methylhistidine (Figure 3D). Taken together, replicative senescence is associated with significant metabolomic changes, which are similar in MSCs and iMSCs.

We have exemplarily validate the senescence-associated changes of orotic acid in cell pellets of MSCs (n = 4) using a fluorogenic reaction (Yin et al., 2015). The results validated a significant increase of orotic acid over subsequent passages (Figure 3E). Thus, the quantification of specific metabolites can be used to track cellular aging.

### The bioenergetic switch in senescence

To gain better insight into how the senescence-associated metabolites map to specific metabolic pathways, we considered all metabolites that were at least significantly changed in MSCs and/or iMSCs. MetaboAnalyst (Xia, Sinelnikov, Han, & Wishart, 2015) indicated that the 213 down-regulated metabolites were particularly associated with glycerophospholipid pathway (p = 2.6 * 10^−7^ and pathway impact score [PI] = 0.47), tricarboxylic acid cycle (p = 9.2 * 10^−6^ and PI = 0.27) as well as taurine and hypotaurine pathways (p = 0.0084 and PI = 0.52; Figure 4A). On the other hand, the 41 up-regulated metabolites were enriched in the biosynthesis of valine, leucine and isoleucine (p = 0.0019 and PI = 0.14), butanoate metabolism (p = 0.0060 and PI = 0.11), pyruvate metabolism (p = 0.0083 and PI = 0.017) and glycolysis (p = 0.015 and PI = 0.18; Figure 4B). These results indicated that changes in tricarboxylic acid cycle and glycolysis might be of particular relevance during senescence, albeit these pathways seemed to be hardly regulated on gene expression level (Supplemental Figure 2).

**Figure 4:**
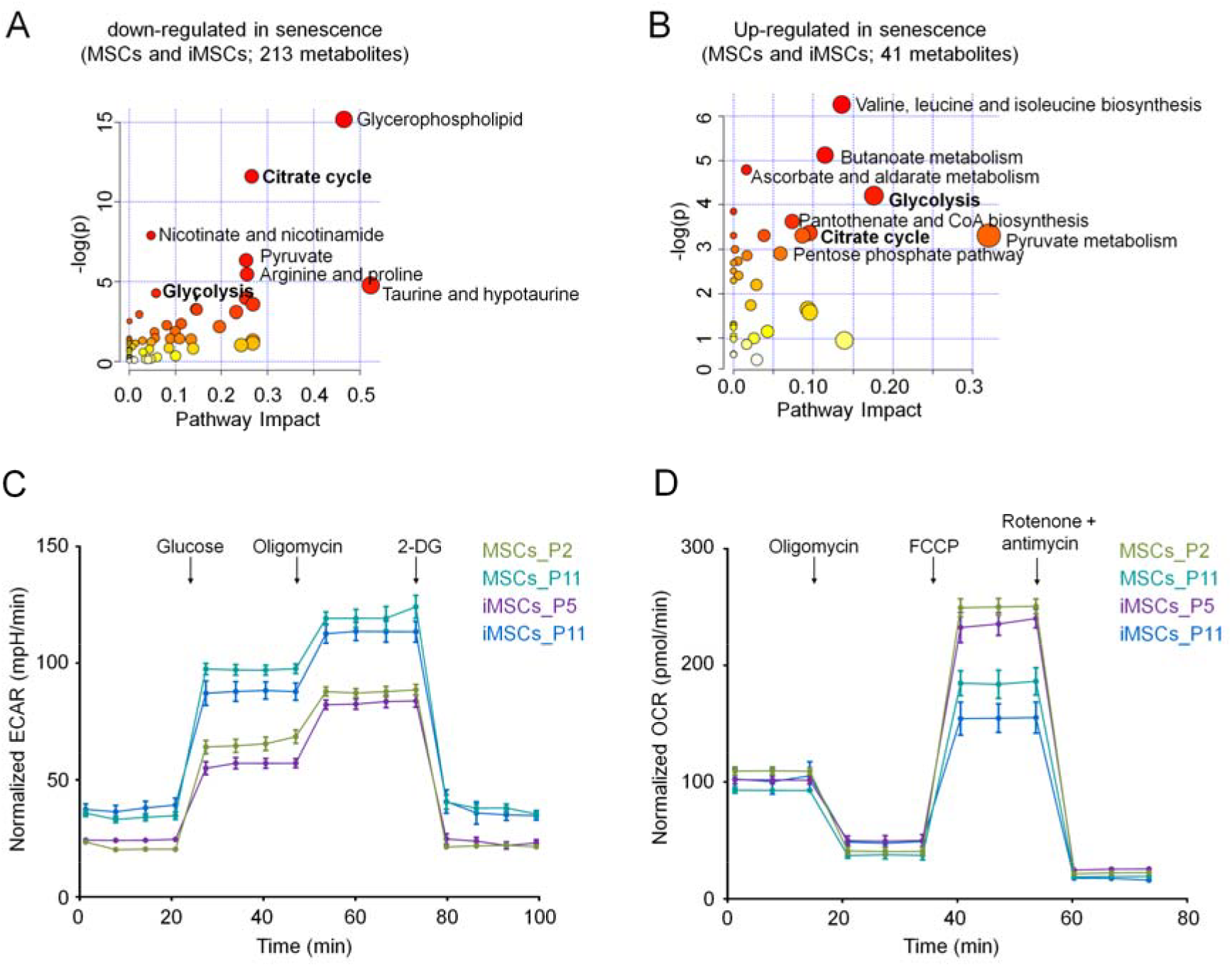
Metabolic pathway analysis during replicative senescence. (A) The metabolic pathway analysis (using MetaboAnalyst 4.0) indicated, that the 213 metabolites, which were significantly down-regulated during senescence of MSCs and/or iMSCs, points towards more oxidative metabolism, whereas (B) the 41 up-regulated metabolites were rather related to glycolytic metabolism. Scores for enrichment (vertical axis) and topology analyses (pathway impact, horizontal axis) are depicted (color code depicts overall significance, and the size of the circles reflects centrality of the involved metabolites). (C) Metabolic flux analysis (Seahorse Bioscience) of the extracellular acidification rate (ECAR) indicates that senescent MSCs and iMSCs use more glycolysis, while (D) the oxygen consumption (OCR) indicates higher oxidative metabolism in MSCs and iMSCs of early passages (n = 8 for each condition).

In fact, previous studies indicated that various cell types gradually shift toward a more glycolytic state during culture expansion (Bittles & Harper, 1984; Zwerschke et al., 2003). We therefore analyzed if this metabolic switch occurs also in MSCs and iMSCs using a flux analyzer to monitor the extra cellular acidification rate (ECAR) and the mitochondrial oxygen consumption rate (OCR). ECAR rather reflects glycolytic behavior and it was consistently higher in senescent MSCs and iMSCs (Figure 4C). In contrast, OCR was higher and more affected in early passages of MSCs and iMSCs (Figure 4D). Thus, both cell types reveal the same senescence-associated shift to glycolytic pathways upon senescence.

## Discussion

Cellular senescence is a very complex stress response that interlinks cell intrinsic and extrinsic processes (Wiley & Campisi, 2016). The results of this study indicate that upon exit from pluripotent state the iPSC-derived MSCs undergo the same functional and molecular changes as previously described for primary MSCs: iMSCs recapitulate senescence-associated changes in morphology, loss of differentiation potential, and ultimately growth arrest. Furthermore, the senescence markers SA-β-gal and the Epigenetic Senescence Signature increased with serial passaging. It might have been anticipated that iMSCs, which are derived from entirely rejuvenated iPSCs, can reach a higher number of cumulative population doublings. For example, MSCs from postnatal placenta have higher proliferative potential than MSC from adult bone marrow (Ventura Ferreira et al., 2018). However, our results indicate that iMSCs enter senescent state even after less cPDs than primary MSCs, indicating that replicative senescence needs to be taken into account for iPSC-derived cell types. On the other hand, the state of cellular aging may be better controlled in iPSC-derived cells by the possibility of expanding iPSCs in pluripotent state to then generate iMSCs of low passage in relevant cell numbers.

Only few studies investigated metabolomic changes during culture expansion and most of them addressed changes in fibroblasts (James et al., 2015; Zwerschke et al., 2003). Overall, there was a consistent shift from oxidative phosphorylation towards glycolysis and the pentose phosphate pathway (James, Lane, Michalek, Karoly, & Parkinson, 2016; Zwerschke et al., 2003). It has been suggested that this was to avoid further damage by reduction of reactive oxygen species (ROS) (James et al., 2016). In fact, particularly for stress-induced premature senescence, ROS are considered important intermediates contributing to the aging phenotype. On the other hand, such damage should rather be avoided in the proliferative iPSCs and early passages. An alternative explanation might be that metabolic changes are caused by a vicious cycle, where depletion of ATP leads to an adaptive response that increases metabolic imbalance (Zwerschke et al., 2003). The shift toward glycolysis at later passages may appear counterintuitive with regard to the general finding that senescent cells have increased numbers of mitochondria (Kim et al., 2018). This might be due to the finding that mitochondria at later passages are often dysfunctional (Zhang, Menzies, & Auwerx, 2018). Despite the fact that many stem cells exhibit low mitochondrial content and rely on mitochondrial independent glycolytic metabolism for energy, accumulating evidence has implicated the importance of mitochondrial function in stem cell activation, fate decisions and defense against senescence (Zhang et al., 2018).

Our results demonstrate that replicative senescence has major impact on the metabolome with highly reproducible changes across different cell types and biological replica. Lee *et al*. have previously identified eight metabolites with altered expression during senescence of MSCs, including lysophosphatidylcholine and lysophosphatidylethanolamine (Lee et al., 2014), while we describe a multitude of significant changes. The highly consistent changes indicate that the metabolic phenotype can be used as a biomarker to assess the state of cellular aging. This may particularly hold true for metabolites that can be quantified by other means, as exemplarily demonstrated for orotic acid, which can be quantified by a fluorogenic reaction. Orotic acid is generated by the mitochondrial enzyme dihydroorotate dehydrogenase or a cytoplasmic enzyme of pyrimidine synthesis pathway, which may again underpin the importance of mitochondria for cellular aging. The metabolomic changes that are consistently observed across different cell types might provide important puzzle pieces in understanding how cellular aging is regulated. This knowledge may also provide new targets for the development of senolytic drugs in the future (Grezella et al., 2018; Zhu et al., 2015).

## Experimental Procedures

### Cell culture of mesenchymal stromal cells

Mesenchymal stromal cells were isolated from the bone marrow of five donors (ranging from 50 to 74 years) after orthopedic hip replacement surgery. All samples were taken after written consent according to the guidelines of the local ethics committees (RWTH Aachen; EK300/13) and isolated as described before (Koch et al., 2013). In brief, cells were flushed from the bone and cultured in parallel in basal medium consisting of Dulbecco’s Modified Eagle Medium (DMEM, 1g/L glucose; PAA, Pasching, Austria) with 1% penicillin/streptomycin (PAA) and 1% L-glutamine, and supplemented with 10% human platelet lysate (HPL). HPL-pools consisted of at least 5 lysates to reduce variation and coagulation was prevented by 0.61 IU unfractionated heparin (UFH; Ratiopharm, Ulm, Germany).

### Generation of iPSCs and differentiation towards MSCs

Induced pluripotent stem cells were reprogrammed from the same MSC donors that were used for experiments with primary MSCs. Reprogramming was performed with episomal plasmids (Willmann et al., 2013). iPSCs were cultured on tissue culture plastic coated with vitronectin (0.5 µg/cm²) in StemMACS iPS-Brew XFe96 (all Miltenyi Biotec GmbH, Bergisch Gladbach, Germany). Pluripotency was validated by *in vitro* differentiation and Epi-Pluri-Score (Cygenia GmbH, Aachen, Germany) (Lenz et al., 2015). iPSCs were re-differentiated toward iMSCs under MSC culture conditions as described above (Frobel et al., 2014). After one week, differentiated iPSCs were maintained on 0.1% gelatine-coated TCP or HPL-gel and passaged every further week using trypsin-EDTA 0.25% (Gibco/Thermo Fisher Scientific).

### Flow Cytometry

Immunophenotypic surface marker analysis was performed on a FACS Canto II (BD, Heidelberg, Germany) upon staining with CD14 allophycocyanin (APC, clone M5E2; BD), CD29 phycoerythrin (PE, clone MAR4; BD), CD31 PE (clone WM59; BD), CD34 APC (clone 8G12; BD), CD45 APC (clone HI30; BD), CD73 PE (clone AD2; BD), CD90 APC (clone 5E10; BD), CD105 fluorescein isothiocyanate (FITC, clone MEM-226; ImmunoTools, Friesoythe, Germany).

### Analysis of Proliferation

At every passage cells were harvested by trypsinization, counted in a Neubauer cell chamber and seeded in defined numbers. Cumulative population doublings (cPDs) were calculated as described before (Cholewa et al., 2011).

### *In vitro* differentiation of MSCs and iMSCs

Differentiation of MSCs and iMSCs toward adipogenic, osteogenic, and chrondrogenic lineage was induced as previously described (Koch et al., 2011). After three weeks, osteogenic differentiation of MSCs and iMSCs was analyzed by staining of calcium precipitates with Alizarin Red S and quantified on a Tecan Infinite 200 plate reader (Gregory, Gunn, Peister, & Prockop, 2004). Fat droplets in adipogenic differentiation were stained with BODIPY counterstained with DAPI. Glycosaminoglycan deposition in chondrogenic differentiation was analyzed by Alcian Blue and PAS staining.

### Transcriptomics

RNA was isolated from the samples using the NucleoSpin RNA extraction kit (Macherey-Nagel, Düren, Germany). Library preparation was performed with the Illumina TruSeq Stranded Total RNA Sample Preparation Kit, with a total input amount of 1 µg, following the standard protocol with ribosomal depletion (Illumina Ribo-Zero Gold rRNA removal kit) and the NextSeq 500/550 High Output Kit v2 (carried out by the Core Genomics-facility IZKF, RWTH Aachen University). The resulting pair-end reads were quality-checked with FastQC (http://www.bioinformatics.babraham.ac.uk/projects/fastqc/), and low-quality reads were removed using TrimGalore. Reads were mapped to the GRCh38 assembly of the human genome using Tophat2 (Trapnell, Pachter, & Salzberg, 2009), version 2.0.10, and reassembled with Cufflinks (Trapnell et al., 2012), version 2.1.1. Differential gene expression was analyzed using the DESeq2 package (Love, Huber, & Anders, 2014), version 1.10.1. Significance of transcriptomic changes was estimated using the R limma package with an adjusted paired p-value <0.05. Gene Ontology analysis was performed with GoMiner™.

### Metabolomics

Pellets of 3 * 10^6^ cells were subjected to methanol extraction, then split into aliquots for analysis by Ultra High Performance Liquid chromatography/Mass Spectrometry/Mass Spectrometry (UHPLC/MS/MS) and Gas Chromatography/Mass Spectrometry (GC/MS) (Tufi et al., 2014). Metabolomics analysis was conducted at Metabolon Inc. (Durham, NC, USA) as previously described (Long et al., 2017). Briefly, Metabolites were identified by automated comparison of ion features to a reference library of chemical standards followed by visual inspection for quality control. Missing values were assumed to be below the detection limits and imputed with the compound minimum (minimum value imputation). To identify the most relevant metabolic pathways during replicative senescence we employed the pathway enrichment analysis for the up- and down-regulated metabolites in MSCs and iMSCs using the pathway analysis tool from MetaboAnalyst 4.0 (www.metaboanalyst.ca). The pathway enrichment analysis used GlobalTest to analyses analyze the concentration values with high sensitivity and to identify subtle changes involved in the same biological pathway (Chong et al., 2018). Statistical tests were performed using R bioconductor package and significance was estimated using one-way ANOVA and adjusted p-value <0.05.

### Quantification of orotic acid

A confluent well of a six-well plate was used to isolate the metabolites with methanol (Ser, Liu, Tang, & Locasale, 2015), while another well was used to determine the total protein concentration to normalize the results. Briefly, 50 μL of the sample was mixed with 50 μL of 4.0 mM of 4-TFMBAO, 50 μL of 8.0 mM K_3_[Fe(CN)_6_] and 50 μL of 80 mM K_2_CO_3_. The mixture (200 μL) was then heated at 80 °C for 4 min, followed by cooling in an ice bath for approx. 2 min to stop the reaction. The relative fluorescence intensity produced by the reaction with 4-TFMBAO was measured with TECAN 200 plate reader (Tecan Group Ltd., Switzerland) at maximum excitation and emission wavelengths of 340 and 460 nm, respectively.

### Metabolic flux analysis

The XFe96 Flux analyzer and Prep Station (Seahorse Bioscience XFe96 Instrument, Agilent, CA, USA) were used according to the manufacturer's instructions (Ferrick, Neilson, & Beeson, 2008). Briefly, cells were seeded in XFe96 cell culture plates at 25,000 cells per well and cultured overnight. The XFe96 sensor cartridges were hydrated overnight with 200 μL of Seahorse Bioscience XFe96 Calibrant at pH 7.4 and stored at 37 °C without CO_2_. One hour before the measurement, cells were washed and culture media were replaced with no-glucose media. Basal measurements of ECAR and OCR, as well as measurements after addition of glucose (final concentration 10 mM; Sigma Aldrich, St. Louis, MO, USA), oligomycin (final concentration 5 μM; Sigma Aldrich) and 2-deoxy-d-glucose (2DG, final concentration 100 mM; Seahorse Bioscience), were performed as described in the XF Glycolysis Stress Test Kit User Manual (Seahorse Bioscience).

## Acknowledgments

This work was funded by the START-Program (to EF-R) and by the Interdisciplinary Center for Clinical Research (IZKF; O3-3; to WW), both within the faculty of Medicine at the RWTH Aachen University; by the Else Kröner-Fresenius-Stiftung (2014_A193; to WW); and by the Deutsche Forschungsgemeinschaft (DFG; WA 1706/8-1 and WA1706/11-1; to WW).

## Author contributions

EF-R and WW developed the study concept and experimental design. EF-R, JF, JH, AO, TS and MO performed the experiments and interpreted the data. J-WK contributed vital reagents and materials, and provided important intellectual support throughout the study. EF-R and WW wrote the first draft of the manuscript. All authors read and approved the final manuscript.

