## Supplemental Figures for "Senescence-associated metabolomic phenotype in primary and iPSC-derived mesenchymal stromal cells"

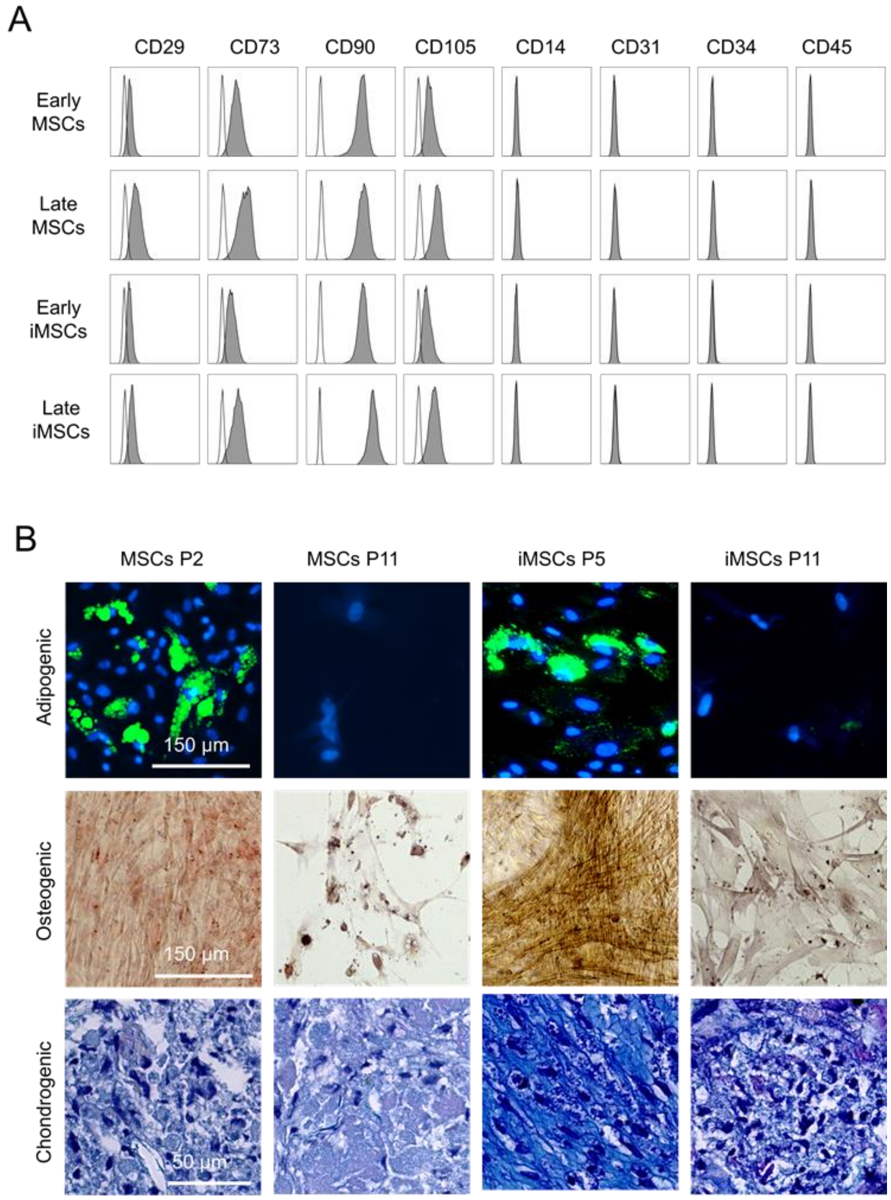


***Figure S1: Immunophenotype and differentiation potential of MSCs and iMSCs.***

*(A) Histograms depict exemplarily the immunophenotype of MSCs and iMSCs in early and late passages. (B) MSCs and iMSCs in early and late passages were differentiated towards adipogenic, osteogenic and chondrogenic lineages and analyzed as indicated in the text.*

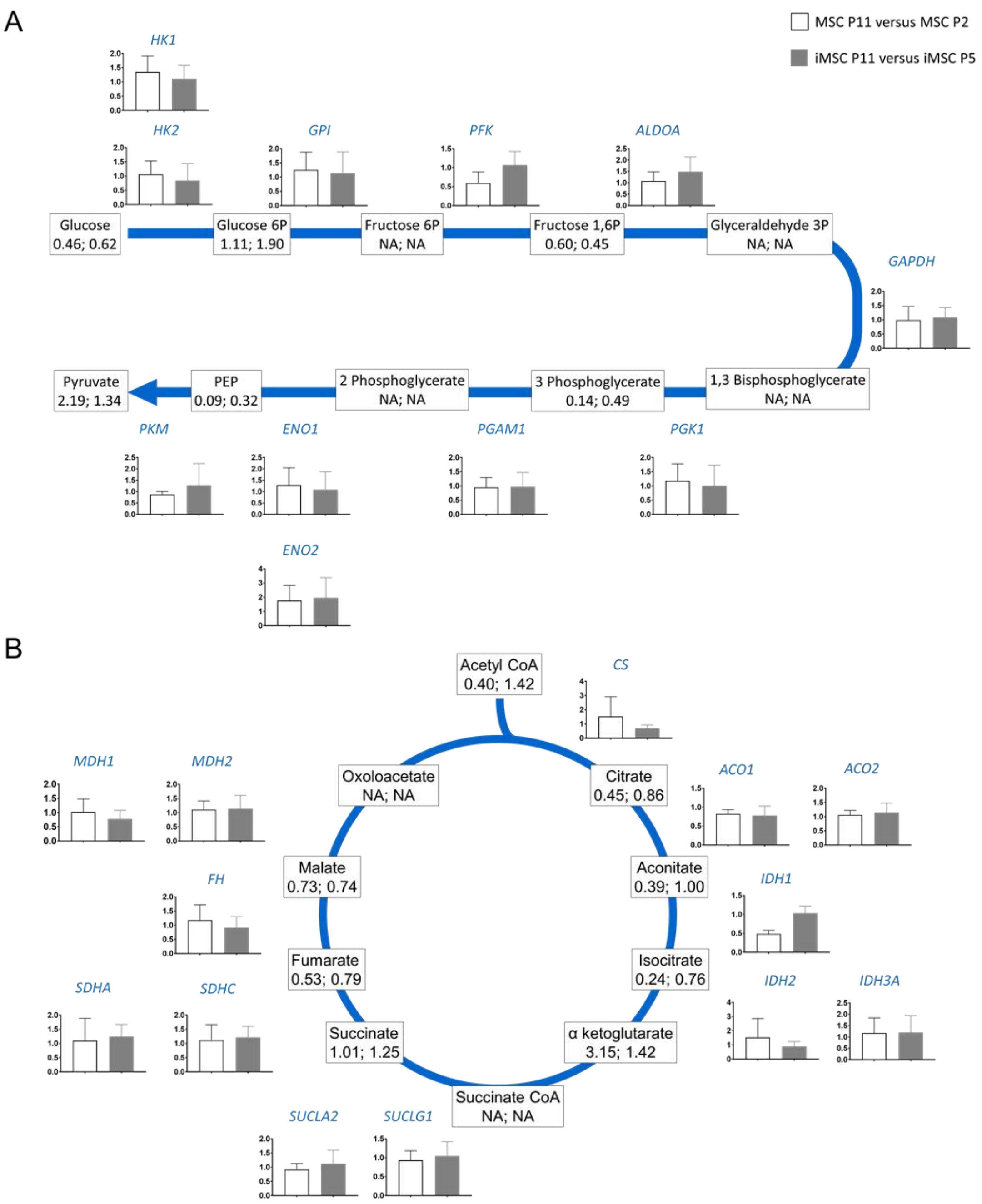


***Figure S2: Metabolomic and transcriptomic changes in glycolysis and tricarboxylic acid cycle.*** *Senescence-associated gene expression changes of enzymes involved in glycolysis (A) and TCA cycle (B) are represented by bars (MSCs P11 versus MSCs P2 in white; and iMSCs P11 versus iMSCs P5 in grey). In addition, the corresponding fold-changes of metabolites are indicated in the boxes (for MSCs and iMSCs, respectively). Abbreviations: HK, hexokinase; GPI, glucose-6-phosphate isomerase; PFK, phosphofructokinase; ALDOA, aldolase A; GAPDH, glyceraldehyde 3-phosphate dehydrogenase; PGK1, phosphoglycerate kinase; PGAM1, phosphoglycerate mutase 1; ENO enolase; PKM, pyruvate kinase; CS, citrate synthase; ACO, aconitase; FH, fumarate hydratase; IDH, isocitrate dehydrogenase; MDH, malate dehydrogenase; SUCL, succinate/malate CoA ligase; SDH, succinate dehydrogenase; NA, not available.*
